# Development of two mouse strains conditionally expressing bright luciferases with distinct emission spectra as new tools for in vivo imaging

**DOI:** 10.1101/2023.04.28.538499

**Authors:** Toshiaki Nakashiba, Katsunori Ogoh, Satoshi Iwano, Takashi Sugiyama, Saori Mizuno-Iijima, Kenichi Nakashima, Seiya Mizuno, Fumihiro Sugiyama, Atsushi Yoshiki, Atsushi Miyawaki, Kuniya Abe

**Author notes:** Correspondence, Contact Information: Toshiaki Nakashiba, Experimental Animal Division, RIKEN BioResource Research Center, Tsukuba, Ibaraki, Japan. Institute for Tenure Track Promotion, University of Miyazaki, Miyazaki, Japan (Satoshi Iwano). R&D division, Evident Corporation, Hachioji, Tokyo, Japan (Takashi Sugiyama).

## Abstract

In vivo bioluminescence imaging (BLI) has been an invaluable noninvasive method to visualize molecular and cellular behaviors in laboratory animals. Bioluminescent reporter mice possessing luciferases for general use have been limited to a classical luciferase, Luc2, from *Photinus pyralis*, and have been extremely powerful for various in vivo studies. However, applicability of reporter mice for in vivo BLI could be further accelerated by increasing light intensity using other luciferases and/or improving the biodistribution of their substrates in animal body. Here, we created two Cre-dependent reporter mice incorporating luciferases: oFluc derived from *Pyrocoeli matsumurai* and Akaluc, both of which had been reported previously to be brighter than Luc2 when using appropriate substrates; we then tested their bioluminescence in neural tissues and other organs in living mice. When expressed throughout the body, both luciferases emitted an intense yellow (oFluc) or far-red (Akaluc) light easily visible to the naked eye. Moreover, oFluc and Akaluc were similarly bright in the pancreas for in vivo BLI. However, Akaluc was superior to oFluc for brain imaging, because its substrate, AkaLumine-HCl, was distributed to the brain more efficiently than the oFluc substrate, D-luciferin. We also demonstrated that the light produced by oFluc and Akaluc was sufficiently spectrally distinct for dual-color imaging in a single living mouse. Taken together, these novel bioluminescent reporter mice are an ideal source of cells with bright bioluminescence and may facilitate the in vivo BLI of various tissues/organs for preclinical and biomedical research in combination with a wide variety of Cre-driver mice.

## Introduction

Bioimaging constitutes an essential part of research in the life sciences, as it provides a wide array of tools for analyses of molecular and cellular behaviors in various biological systems. Since the advent of fluorescent protein (FP) technology, live imaging has revolutionized the fields of cell biology because in theory, any gene product or type of cell can be marked and traced ^1^. However, it has been challenging to apply FP technology to live imaging at the organismal level. The excitation light required for FP technology in biological tissues is severely attenuated because of light scattering and absorption, rendering in vivo live imaging using FP impractical ^2, 3^. Furthermore, the excitation light causes “autofluorescence” from biological materials, such as NADPH and flavoproteins, which exacerbates the signal-to-noise ratio of FP signals ^4, 5^.

By contrast, bioluminescence imaging (BLI) using bioluminescent luciferase/luciferin systems has a wider dynamic range than fluorescent imaging because of the lack of requirements for excitation light ^6, 7^. BLI is considered to be a more suitable modality for noninvasive deep-tissue imaging. Among the bioluminescent luciferase/luciferin systems, the luciferase from the firefly, *Photinus pyralis*, particularly its derivative Luc2, has been the most widely used reporter; it produces an orange light by oxidizing D-luciferin (peak emission at 609 nm) ^8^. Although reporter mice possessing Luc2 have been widely used for in vivo BLI ^9–12^, there remains some issues, e.g. relatively weak light intensity produced from the luciferase and the limited biodistribution of D-luciferin, which distributes poorly within the brain because of the blood‒brain barrier (BBB) ^13^.

For better in vivo BLI, attempts have been made to increase the light intensity and improve the biodistribution of substrates. As synthetic compounds of D-luciferin derivative, CycLuc1, and AkaLumine hydrochloride (AkaLumine-HCl) exhibit enhanced biodistribution in most tissues including brain and produce red-shifted light, which could result in a better penetration of animal tissues and bodies (the peak emission of Luc2 is at 604 nm for CycLuc1 and at 677 nm for AkaLumine-HCl) ^14, 15^. Meanwhile, novel luciferase derivatives with a brighter light intensity and/or red-shifted wavelength have been searched ^16–18^. Although a red-shifted wavelength was achieved by site-directed mutagenesis of Luc2 using D-luciferin as a substrate, the total photon yield in most of these mutants did not exceed that produced by the original Luc2 ^17^. This poor yield was also the case when combining Luc2 and the synthetic luciferins mentioned above ^19^. More recently, Iwano et al. ^20^ reported the AkaBLI system, in which an innovative derivative of Luc2, Akaluc, was used in conjunction with AkaLumine-HCl. Remarkably, Akaluc produced a luminescent signal that was 10 times brighter than that produced by Luc2 when measured in vitro. Even more remarkably, the difference exceeded 100-fold for in vivo BLI, probably because of the emission peak of AkaBLI at 650 nm, which is within the range of the “optical window of biological tissues” ^4^. In fact, single cells trapped in the mouse lung could be detected using the AkaBLI system ^20^. This newly developed BLI system should open new avenues of research involving deep-tissue imaging.

Newly developed BLI systems have been mostly evaluated in in vivo settings by injecting cells or viral vectors carrying luciferase genes into small animals ^14, 15, 17, 19–22^. However, these methods are limited to cells that are amenable to viral transduction and to several spatial locations in the body that are accessible using procedures such as the pulmonary trapping of intravenously injected cells or subcutaneous cellular transplants. Furthermore, surgical procedures for viral injection or cellular transplants may have various consequences in different animals when considering the number of cells and their spatial location, thus complicating the evaluation of BLI systems. Therefore, the next logical step in the development of the BLI system is the generation of genetically modified mouse strains in which the BLI system can be systematically operated by genetic means.

Another item to add to the BLI toolbox would be bright luciferases with an emission peak distinct from that of Akaluc. Multicolor imaging is critical and advantageous for bioimaging to detect and analyze the behaviors or interactions of more than two elements in biological systems (as exemplified by multicolor imaging using multiple FPs with distinct emission peaks). However, for in vivo BLI, luciferases that are as bright as Akaluc but exhibit different emission peaks have not yet been reported. Ogoh et al. ^23^ isolated and characterized a luciferase derived from the firefly *Pyrocoeli matsumurai*, which inhabits Okinawa Island of Japan. This luciferase (hereinafter referred to as oFluc) produces yellow light (peak emission at 567 nm) when D-luciferin is used as a substrate, and its luminescence intensity is 10 times brighter than that produced by Luc2 when measured in cultured cells ^23^. While Akaluc was found not to catalyze D-luciferin for light production ^20^, the substrate specificity of oFluc has not yet been clarified. Based on the assumption that oFluc does not produce light with AkaLumine-HCl, we envisaged that oFluc might be an ideal partner for Akaluc for dual-color BLI using both substrates.

In the present study, we generated two mouse strains, in which Cre-dependent reporter constructs carrying Akaluc and oFluc were inserted into the *ROSA26* locus, which is a safe harbor site in the mouse genome for stable expression ^24^. By crossing with a general-deleter Cre strain, we created “glowing mice” that exhibited whole-body bioluminescence, thus representing an ideal source of various bioluminescence-emitting cells to suit a wide variety of transplantation studies. These mouse strains can certainly be directed to express the luciferase reporters in specific cell populations using the Cre/loxP recombination system, thereby allowing the noninvasive imaging of specific cell populations in the whole body. Furthermore, this system should be helpful in locating cell domains undergoing Cre-mediated recombination during development and adulthood. Although such Cre-mediated expression of Luc2 in the mouse has been reported ^25–28^, the use of much brighter BLI systems, such as Akaluc and oFluc, with distinct emission peaks will undoubtedly expand the utility of the reporter lines for both basic and preclinical research. Here, we demonstrated the feasibility of in vivo dual-color BLI in a single living mouse that harbored both the Akaluc and oFluc reporters.

## Results

### Generation of oFluc and Akaluc reporter mice

We aimed to introduce reporter constructs (Figure 1a) carrying oFluc or Akaluc into the *ROSA26* locus, a safe-harbor for stable expression in the mouse genome. The expression of the luciferases was driven by the strong and ubiquitous *CAG* promoter ^29^. A floxed transcription stop cassette̶the neomycin resistance gene (*Neo*) followed by SV40 poly A̶was placed between the *CAG* promoter and the reporter. The stability of the reporter mRNA was further enhanced by the woodchuck hepatitis virus posttranscriptional regulatory element (WPRE) ^30^. The Akaluc-encoding cDNA was fused at its N-terminus with Venus (Venus/Akaluc) to compare BLI and fluorescent imaging via equimolar expression of the two reporters. By contrast, oFluc was not fused to maximize its expression (Figure 1a).

**Figure 1.**
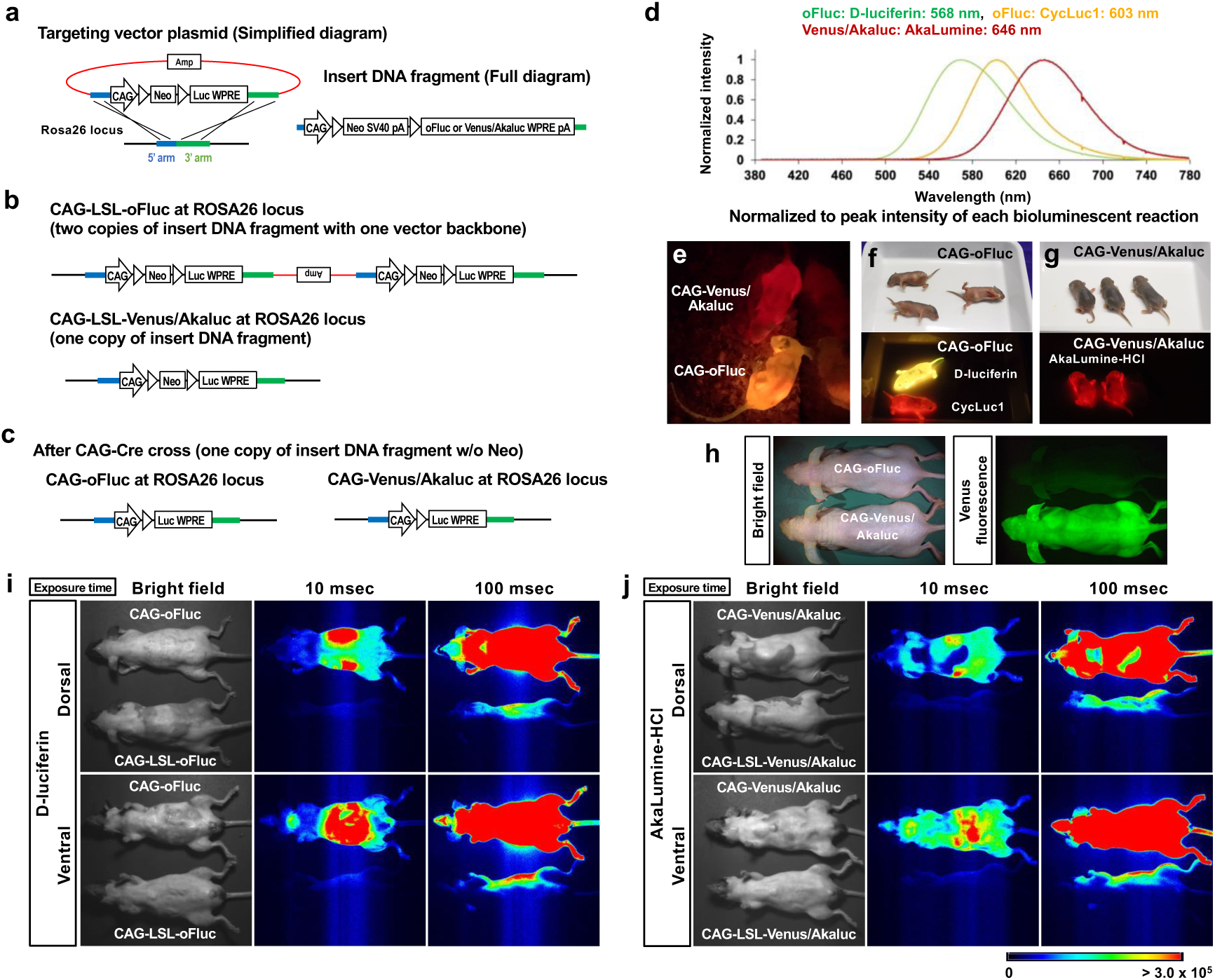
Luciferase reporter mice and in vivo BLI of “glowing mice” (a) Simplified diagram of the targeting vector plasmid (left) for the *ROSA26* locus and a full description of the insert DNA fragment (right). The plasmid backbone, 5′ homology arm, and 3′ homology arm are indicated in red, blue, and green, respectively. (b) Cre-dependent luciferase knock-in alleles at the *ROSA26* locus; two copies of the CAG-LSL-oFluc insert DNA fragment were integrated together with a plasmid backbone, whereas one copy of the CAG-LSL-Venus/Akaluc insert DNA fragment was integrated without a plasmid backbone. (c) After the CAG-Cre cross, both CAG-oFluc and CAG-Venus/Akaluc mice had one copy of the insert DNA fragment without the *Neo* cassette. (d) Emission spectra of luciferases with their substrates. (e) Representative image of CAG-oFluc and CAG-Venus/Akaluc mice captured using a digital color camera at 10 min after IP injection of their substrates (100 mM D-luciferin, 5 μl/gbw for CAG-oFluc; 15 mM AkaLumine-HCl, 5 μl/gbw for CAG-Venus/Akaluc). Freely behaving mice were recorded at 30 frames per second (see Supplementary Movie 1). (f and g) Representative images of mice at postnatal day 4 were captured using a digital color camera after IP injection of the substrate (10 μl). The mice on the right side received no substrate injection. (h) Venus fluorescence of CAG-Venus/Akaluc mice. (i and j) In vivo BLI images were captured using an EM-CCD camera. CAG-oFluc was compared with the negative control, CAG-LSL-oFluc (i), and CAG-Venus/Akaluc was compared with CAG-LSL-Venus/Akaluc (j), (100 mM D-luciferin, 5 μl/gbw for oFluc; 15 mM AkaLumine-HCl, 5 μl /gbw for Venus/Akaluc). The hair on both the dorsal and ventral sides of the mice was shaved. The signals from the bodies of CAG-LSL-oFluc and CAG-LSL-Venus/Akaluc mice reflected the light emitted from the CAG-oFluc and CAG-Venus/Akaluc mice placed next to these animals (note that the signals were found only on the near side of the bodies).

We generated two mouse strains, CAG-LSL-oFluc and CAG-LSL-Venus/Akaluc, using the CRISPR‒Cas9 technique with fertilized eggs collected from C57BL/6J mice. Both strains integrated the insert DNA at the *ROSA26* locus (Figure 1b and Supplementary Figure S1). One copy of CAG-LSL-Venus/Akaluc was inserted into the locus, whereas two copies of CAG-LSL-oFluc were inserted with a vector backbone of the targeting vector plasmid. After crossing these animals with CAG-Cre mice ^31^, for germ-line recombination, we established two additional mouse strains, CAG-oFluc and CAG-Venus/Akaluc, both of which carried one copy of the insert DNA fragment from which the stop cassette was deleted from the *ROSA26* locus (Figure 1c and Supplementary Figure S1).

### “Glowing mice” in two different colors

We examined the bioluminescence of CAG-oFluc and CAG-Venus/Akaluc mice in a dark room. We intraperitoneally (IP) injected 100 mM D-luciferin (5 μl/gbw) into CAG-oFluc mice, and 15 mM AkaLumine-HCl (5 μl/gbw) into CAG-Venus/Akaluc mice. Shortly after the IP injection into live animals, mice of both strains started emitting light, initially from their abdomen and, after 5 min, throughout their body. These signals were highly bright and even visible to the naked eye. The movements of live mice could be recorded by a consumer-grade digital color camera (Sony α7SII) at the video-frame rate (of 30 frames per second (fps)) (Figure 1e–g, Supplementary Movie 1). Both luciferases were intensely bright in the BLI setup using an EM-CCD camera. Their signals saturated the EM-CCD sensor with an exposure of only tens of milliseconds (Figure 1i and j). By contrast, the negative controls showed no detectable signal during the same exposure time. Signals from post-implantation embryos comprising approximately 1000 cells ^32^ and carrying the reporters were successfully detected in utero at embryonic day 6.5 (E6.5) (Supplementary Figure S2). CAG-oFluc and CAG-Venus/Akaluc mice emitted light with comparable intensity in distinct colors corresponding to their peak emissions, yellow and red, respectively (Figure 1d–g). The injection of a synthetic luciferin, CycLuc1 ^15^, into CAG-oFluc mice produced red-shifted light (Figure 1f), which was consistent with the data obtained in vitro using recombinant luciferase proteins and substrates (Figure 1d). Anesthetized CAG-Venus/Akaluc mice exhibited Venus fluorescence in their entire body (Figure 1h).

### Luciferase activity in tissues and their extracts

We then quantitatively assessed luciferase activity in crude tissue extracts via incubation with their appropriate substrates (Figure 2a–c). Five groups of mice were included in this assay: CAG-oFluc, CAG-Venus/Akaluc, CAG-LSL-oFluc, CAG-LSL-Venus/Akaluc, and C57BL/6. CAG-oFluc and CAG-Venus/Akaluc mice displayed strong bioluminescent signals in all tissues examined; the increase in the signals relative to the negative control (C57BL/6) mice was on average 10^4^- or 10^3^-fold, respectively. By contrast, the signals in CAG-LSL-oFluc and CAG-LSL-Venus/Akaluc mice were extremely low, comparable with those of C57BL/6 mice, indicating the robust inducibility of luciferase expression in a Cre-dependent manner. Of note, these mice exhibited a slight but significant increase in signal strength over C57BL/6 mice in some tissues, such as the brain and testis.

**Figure 2.**
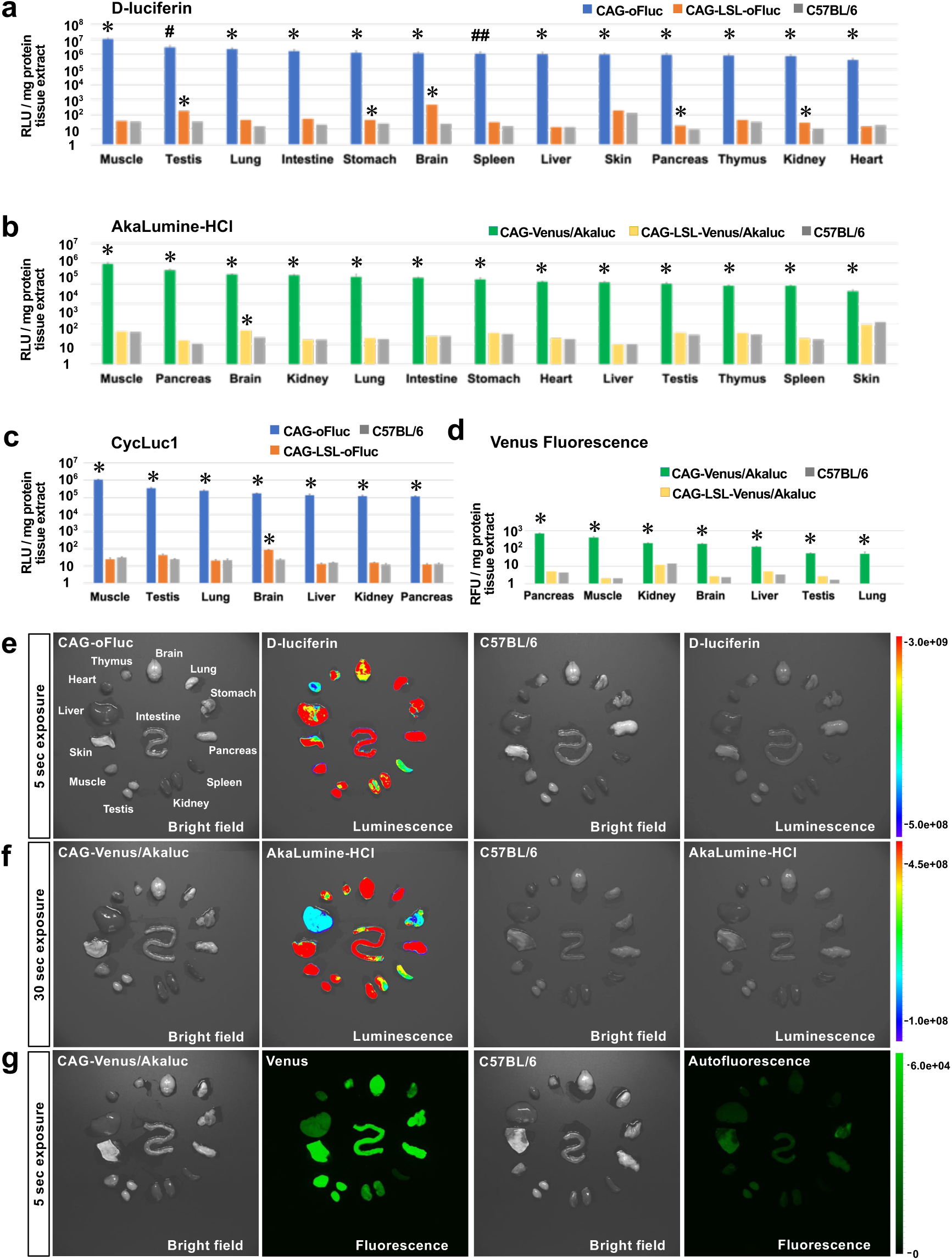
Luciferase activity in tissues and their extracts (a and b) Relative luminescence units (RLU) normalized to mg of protein tissue extracts of CAG-oFluc (a) and CAG-Venus/Akaluc (b) mice supplemented with their substrates (1 mM each, D-luciferin and AkaLumine-HCl, respectively). For comparison, tissue extracts were collected before the CAG-Cre cross, i.e., CAG-LSL-oFluc and CAG-LSL-Venus/Akaluc, and from C57BL/6 mice are included in the graphs. From left to right, tissues exhibiting stronger signals were plotted in order. (c) RLU of the tissue extracts of oFluc supplemented with 1 mM CycLuc1. (d) Relative fluorescence units for the Venus reporter in CAG-Venus/Akaluc. n = 3 mice per group for tissue extracts. ^∗^*P* < 0.05, ^#^*P* = 0.057, and ^##^*P* = 0.059 compared with C57BL/6 mice (*t* test). (e and f) Ex vivo imaging of tissues dissected from CAG-oFluc (e) and CAG-Venus/Akaluc (f) mice incubated with their substrates (1 mM). As negative controls, tissues from C57BL/6 mice were imaged under the same conditions (right). Scale, photon/s/cm^2^/sr. (g) Ex vivo imaging of Venus fluorescence in tissues from CAG-Venus/Akaluc mice. Note that C57BL/6 mice exhibited autofluorescence in some tissues when imaged under the same conditions. Scale, intensity (0–65,025).

The fold increase in the fluorescent intensity of Venus in CAG-Venus/Akaluc mice was not as large as that of Akaluc bioluminescence and was within the range of a 10- to 200-fold increase relative to C57BL/6 mice (Figure 2d). This relatively small fold increase was attributed to the high background signal caused by autofluorescence, whereas no such background was detected in bioluminescence, as revealed by the ex vivo imaging of tissues incubated with their appropriate substrates (Figure 2e–g).

### In vivo BLI of specific cells/organs

The experiments described above were conducted by incubating tissue extracts and dissected organs with the substrates; thus, they did not consider the biodistribution of the substrates and light penetration throughout the mouse bodies. We next directed the expression of reporter luciferases in specific tissues/organs using cell-type-specific or tissue-specific Cre-driver mice for in vivo BLI. We selected four Cre-driver strains that express Cre in neural or nonneural tissues. In vivo BLI was performed using an EM-CCD camera at 10 min after the IP injection of the substrates (100 mM D-luciferin, 5 μl/gbw; 15 mM AkaLumine-HCl, 5 μl/gbw). The signals were captured through a series of exposure times, and raw images are presented together for intuitive assessment in Figures 3 and 4.

**Figure 3.**
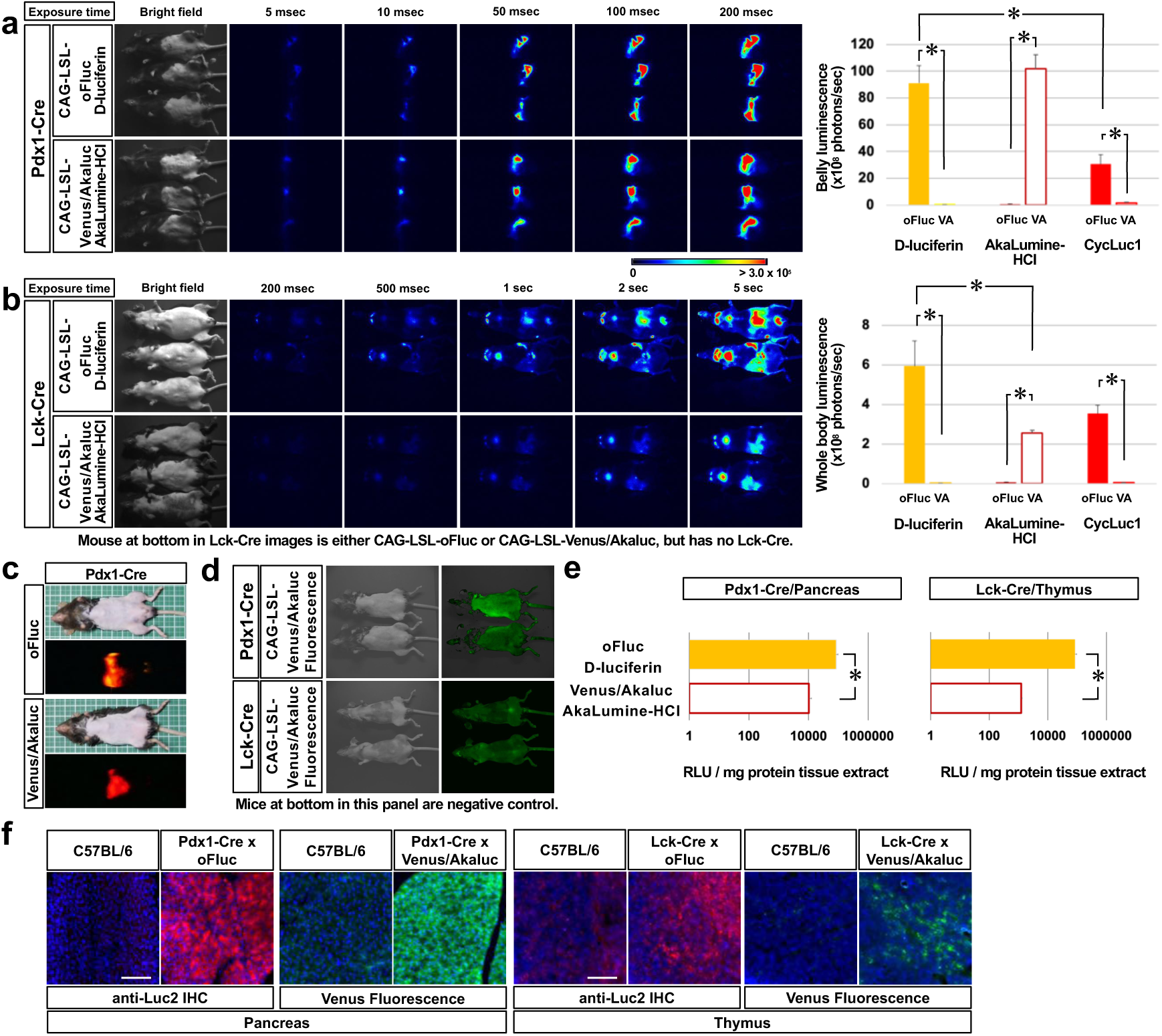
In vivo BLI of nonneural tissues (a) In vivo BLI of oFluc and Akaluc in the pancreas by crossing Pdx1-Cre mice with CAG-LSL-oFluc or CAG-LSL-Venus/Akaluc mice. Images were acquired at multiple exposure times. The hair on the lower abdomen of the mice was shaved. Right panel: quantification of belly luminescence (n = 5 each) for three substrates. Closed boxes, oFluc; open boxes, Akaluc (VA). ^∗^*P* < 0.05 (*t* test). (b) In vivo BLI of oFluc and Akaluc in T cells by crossing Lck-Cre mice with CAG-LSL-oFluc or CAG-LSL-Venus/Akaluc mice. Note that the signal intensity was relatively weak; therefore, Cre-dependent reporter mice before Cre crossing are included at the bottom of each panel (b) as a negative control. The hair on the abdominal area of the mice was shaved. Scale, photon/s/cm^2^. Right panel: quantification of whole-body luminescence (n = 5 each) for three substrates. ^∗^*P* < 0.05 (*t* test). (c) Images captured using a digital color camera. (d) Venus fluorescence from the experimental mice (top) and negative control mice (bottom). Note that experimental mice double positive for Cre and Venus/Akaluc were indistinguishable from negative control mice lacking Venus/Akaluc. (e) RLU of mg protein tissue extracts. ^∗^*P* < 0.05 (*t* test). (f) Histological confirmation of reporter expression in the pancreas or thymus using anti-Luc2 immunohistochemistry (IHC, red color) or native Venus fluorescence (green). Counterstained with DAPI (blue). Scale bar (100 μm).

**Figure 4.**
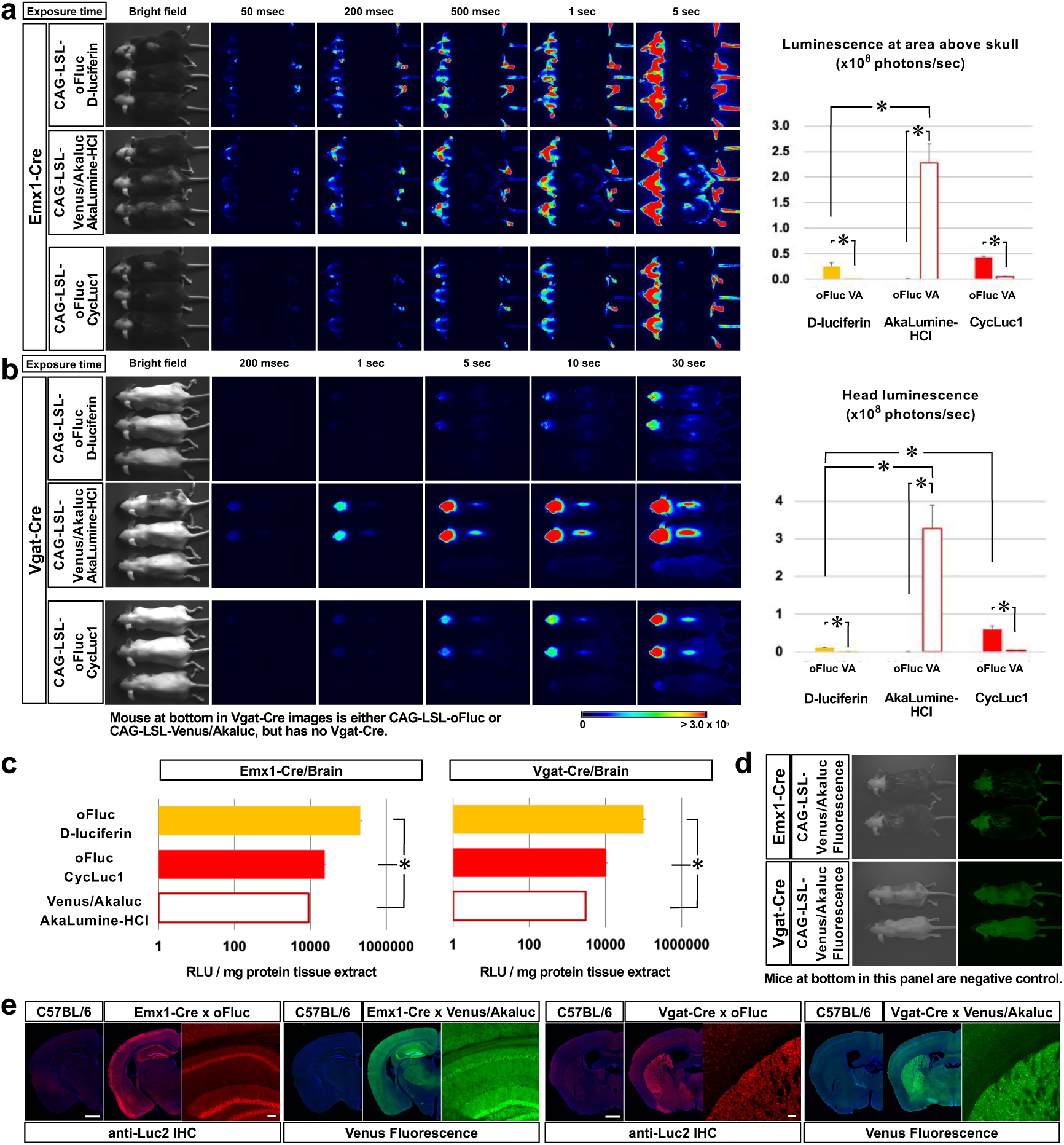
In vivo BLI in neural tissues (a) In vivo BLI for oFluc and Akaluc in mice that were double positive for Emx1-Cre and the reporters. Images were acquired at multiple exposure times. The hair of mice in areas of the head and neck was shaved. Right panel: quantification of luminescence in the area above the skull (n = 5 each) for three substrates. Closed boxes, oFluc; open boxes, Akaluc (VA). ^∗^*P* < 0.05 (*t* test). (b) In vivo BLI using Vgat-Cre . Note that the signal intensity was relatively weak; therefore, Cre-dependent reporter mice before Cre crossing were included at the bottom of each panel (b) as a negative control. The hair on the dorsal side of the mice was shaved. Scale, photon/s/cm^2^. Right panel: quantification of head luminescence (n = 5 each) for three substrates. ^∗^*P* < 0.05 (*t* test). (c) RLU of mg protein tissue extracts. ^∗^*P* < 0.05 (*t* test) in any of the pairs. (d) Venus fluorescence of experimental mice (top) and negative control mice (bottom). The fluorescent signals in experimental mice that were double positive for Cre and Venus/Akaluc were indistinguishable from the autofluorescence obtained in the negative control mice. (e) Histological confirmation of reporter expression in forebrain sections using anti-Luc2 IHC (red) or native Venus fluorescence (green). Counterstained with DAPI (blue). Scale bars, 1 mm for images of whole coronal sections and 100 μm for magnified images).

For the analysis of nonneural tissues, we used Pdx1-Cre and Lck-Cre mice, which have expression specificities for the pancreas/duodenum ^33^ or T cells ^34^, respectively. After crossing with Pdx1-Cre mice (Figure 3a), the oFluc and Akaluc signals were detectable in the upper abdomen after an exposure of only 5 ms, and were saturated at an exposure of 50 ms or longer. The signal intensities of oFluc and Akaluc were comparable with each other and were sufficiently strong to be captured by a consumer-grade color camera; moreover, their signals could be distinguished based on the emission spectra (Figure 3c). After crossing with Lck-Cre mice (Figure 3b), the BLI signals appeared to be restricted to the thymus and other lymphoid organs, and oFluc signals were detectable after an exposure of ∼200 ms and saturated after an exposure of a few seconds. These signals were apparently generated in a Cre-dependent manner (Figure 3b, the animals at the bottom of each panel are CAG-LSL-oFluc or CAG-LSL-Venus/Akaluc mice before Cre cross, respectively, and serve as negative controls). Although oFluc produced a much brighter light than Akaluc in thymus extracts incubated with their substrates (Figure 3e), these differences were greatly attenuated in in vivo BLI, resulting in 2.3-fold stronger signals from oFluc vs. Akaluc (Figure 3b). Histological results confirmed both oFluc and Venus/Akaluc expression in the target tissues (Figure 3f); however, the target-tissue-specific expression of Venus fluorescence was undetectable from outside the body (Figure 3d).

We then crossed the reporter strains with Emx1-Cre or Vgat-Cre (also known as Slc32a1-Cre) mice, to visualize the excitatory neurons of the dorsal forebrain ^35^ or the inhibitory neurons of all neural tissues ^36^ (Figure 4). We confirmed oFluc and Venus/Akaluc expression by measuring enzymatic activities in brain tissue extracts and via histological analysis (Figure 4c and e). After crossing with Emx1-Cre mice (Figure 4a), BLI signals were detected in the head, as expected. However, it was surprising to observe the signals in other sites of the body, such as the base of the ears, hindlimbs, and tail, thus revealing hitherto unknown domains of Cre expression/recombination. After crossing with Vgat-Cre mice (Figure 4b), the BLI signals were confined to the head and spine, with the latter presumably corresponding to inhibitory neurons in the spinal cord. As predicted from the poor biodistribution of D-luciferin in the brain, a longer exposure time was required to detect oFluc signals through the skull. By contrast, Akaluc signals through the skull could be readily detected within 100–200 ms (the head luminescence in Akaluc/AkaLumine-HCl mice was 30-fold greater than that detected in oFluc/D-luciferin mice; Figure 4b, right graph). With CycLuc1 (5 mM, 10 μl/gbw), oFluc signals were greatly increased through the skull (at approximately 5.4-fold over D-luciferin); however, this signal level did not reach to the level achieved by Akaluc (Figure 4b). Thus, Akaluc/AkaLumine-HCl was the best luciferase/luciferin combination for in vivo BLI in brain tissues. None of the Cre crosses yielded Venus fluorescence signals in in vivo imaging (Figure 4d).

### In vivo BLI of CAG-LSL reporter mice prior to the Cre cross

From the results of the in vitro luciferase assay, we noticed that both CAG-LSL-oFluc and CAG-LSL-Venus/Akaluc mice exhibited very low, albeit significant, luciferase activity in some tissues (Figure 2a–c). To confirm this observation using in vivo BLI, we performed BLI in both CAG-LSL reporter mice using an EM-CCD camera after the IP injection of the substrates (100 mM D-luciferin, 5 μl/gbw; 15 mM AkaLumine-HCl, 5 μl/gbw; Figure 5). Using an exposure time of 1 min or longer, oFluc signals appeared around the kidney and testis, whereas Akaluc signals were detected in the head region (Figure 5a and b). C57BL/6 mice showed no signals. These suspected tissues were consistent with those exhibiting a slight but significant luciferase activity in the tissue extracts (Figure 2a and b). This finding was further confirmed by ex vivo imaging of the brain and testis (Figure 5e and Supplementary Figure S3).

**Figure 5.**
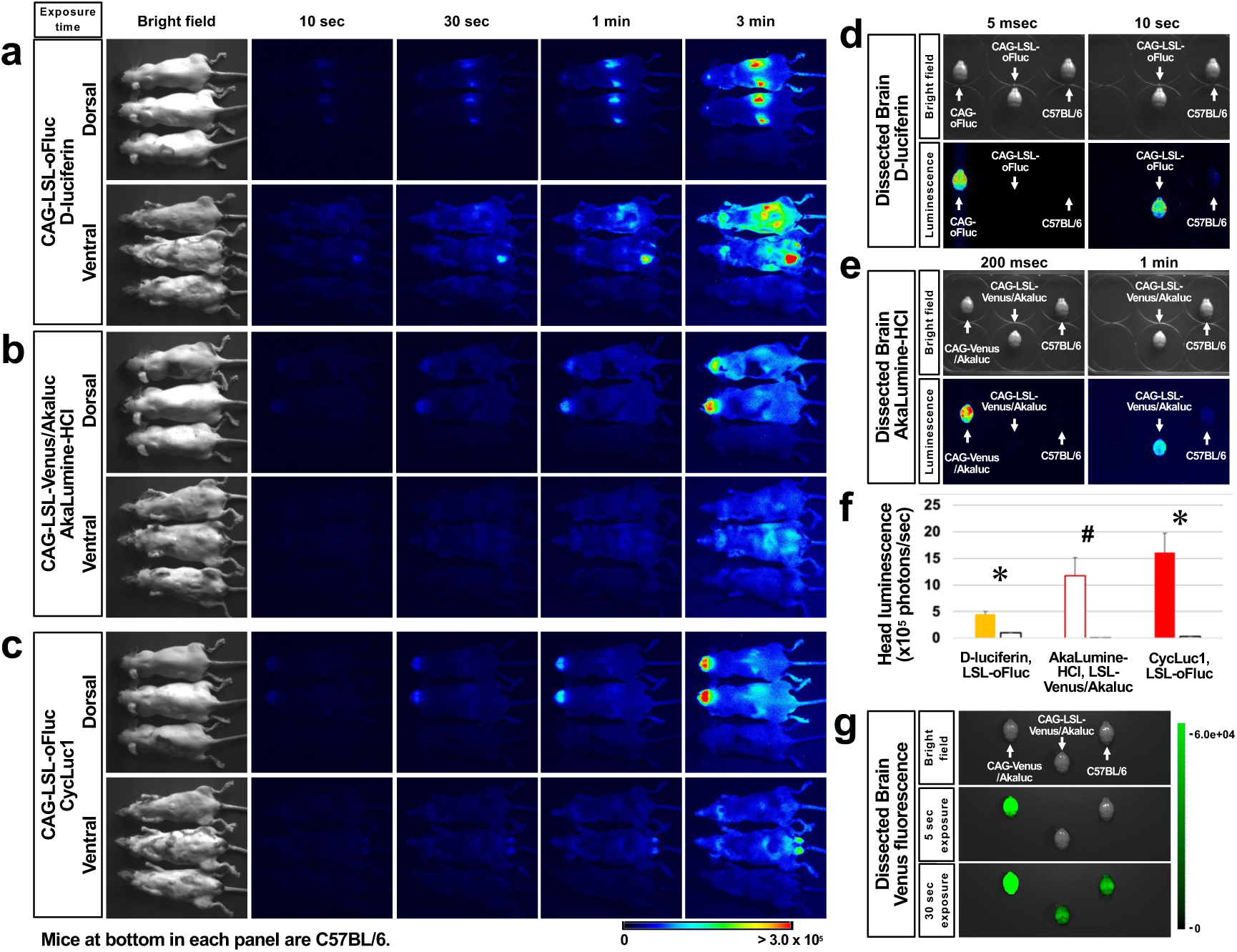
In vivo BLI of Cre-dependent reporter mice before the Cre cross (a and b) Basal luminescence of CAG-LSL-oFluc (a) and CAG-LSL-Venus/Akaluc (b) mice at 10 min (dorsal side) and 50 min (ventral side) after IP injection of their substrate (100 mM D-luciferin, 5 μl/gbw for oFluc; 15 mM AkaLumine-HCl, 5 μl/gbw for Akaluc). Images acquired using four different exposure times are shown, and each panel includes three mice (from the top, two experimental mice and a C57BL/6 mouse, as a negative control; mouse sex: female, male, and male, respectively). The hair on their dorsal and ventral sides of the mice was shaved. (c) Basal luminescence of CAG-LSL-oFluc mice after IP injection of CycLuc1 (5 mM, 10 μl/gbw). Images were acquired using the same mice as shown in (a). Scale, photon/s/cm^2^. (d and e) Ex vivo imaging of dissected brains incubated with D-luciferin (1 mM) or AkaLumine-HCl (1 mM). For longer exposure, the dissected brains of CAG-oFluc or CAG-Venus/Akaluc mice were removed for BLI because their bright luminescence also lit up other tissues, resulting in false-positive signals. (f) Quantification of head luminescence for in vivo BLI. The yellow and red filled boxes correspond to oFluc luminescence using D-luciferin and CycLuc1, respectively. The red open box corresponds to Akaluc luminescence using AkaLumine-HCl. The black open boxes correspond to luminescence in C57BL/6 mice. n = 4 for reporter mice and n = 3 for C57BL/6 mice. ^∗^*P* < 0.05 and ^#^*P* = 0.054 compared with C57BL/6 mice (*t* test). (g) Venus fluorescence of dissected brains.

Although the results of the enzymatic assay in the tissue extracts indicated very low but significant luciferase activity in the brains of CAG-LSL-oFluc mice (Figure 2a), in vivo BLI failed to detect a signal from their heads (Figure 5a). We hypothesized that D-luciferin is not delivered efficiently into the brain in vivo compared with AkaLumine-HCl. In fact, ex vivo imaging of dissected brains incubated with their substrates revealed signals in CAG-LSL-oFluc mice (Figure 5d), similar to CAG-LSL-Venus/Akaluc mice (Figure 5e). Furthermore, CycLuc1 injection (5 mM, 10 μl/gbw) into CAG-LSL-oFluc improved signal detection in the head (Figure 5c and f). Thus, although the Cre-dependent mice were initially designed to prevent luciferase expression by the transcription stop cassette, these results indicated that very weak but significant leaky luciferase expression occurred in some tissues. In this sense, BLI was more sensitive than fluorescent imaging, because leaky expression of Venus could not be detected or distinguished from the high background autofluorescence (Figure 5g).

### In vivo dual-color BLI

We designed in vivo dual-color BLI experiments using oFluc and Akaluc in the hope that their emissions would be practically separable (Figure 1d). We first tested substrate cross-reactivity using recombinant luciferase proteins. We found that both oFluc and Akaluc emitted almost no signal when using their inappropriate substrate (oFluc/AkaLumine and Akaluc/D-luciferin; hereinafter referred to as mismatched pairs) in vitro (Supplementary Figure S4). We next evaluated the cross-reactivity of substrates in the in vivo setting that used CAG-LSL-oFluc and CAG-LSL-Venus/Akaluc mice individually (Supplementary Figures S5–S8). We confirmed that the original luciferase/substrate pairs (oFluc/D-luciferin and Akaluc/AkaLumine-HCl; hereinafter referred to as matched pairs) produced much stronger signals (more than 88- to 394-fold) than the mismatched pairs (oFluc/AkaLumine-HCl and Akaluc/D-luciferin) (bar graphs in Figure 3a and b).

Our final challenge was to achieve in vivo dual-color BLI in single mice. To localize both oFluc and Venus/Akaluc deep in the same subject, we generated female mice carrying two types of “glowing fetuses” (Figure 6). Fertilized eggs carrying CAG-oFluc and CAG-Venus/Akaluc were collected and transferred separately to the left and right sides, respectively, of the uterus of wild-type mice. We conducted in vivo BLI experiments at the late-gestation stage of E14.5, when fetal positions are largely inferable from the outside (Figure 6). However, at this stage, the blood-placenta barrier is established ^37^ and may hinder the intra-fetal biodistribution of D-luciferin. Thus, to redress the concentration balance between the two substrates, we modified the injection doses— doubling the amount of D-luciferin (100 mM, 10 μl/gbw) and reducing the amount of AkaLumine-HCl to 1/5 or 1/10 (3 or 1.5 mM, 5 μl/gbw) relative to the standard doses.

**Figure 6.**
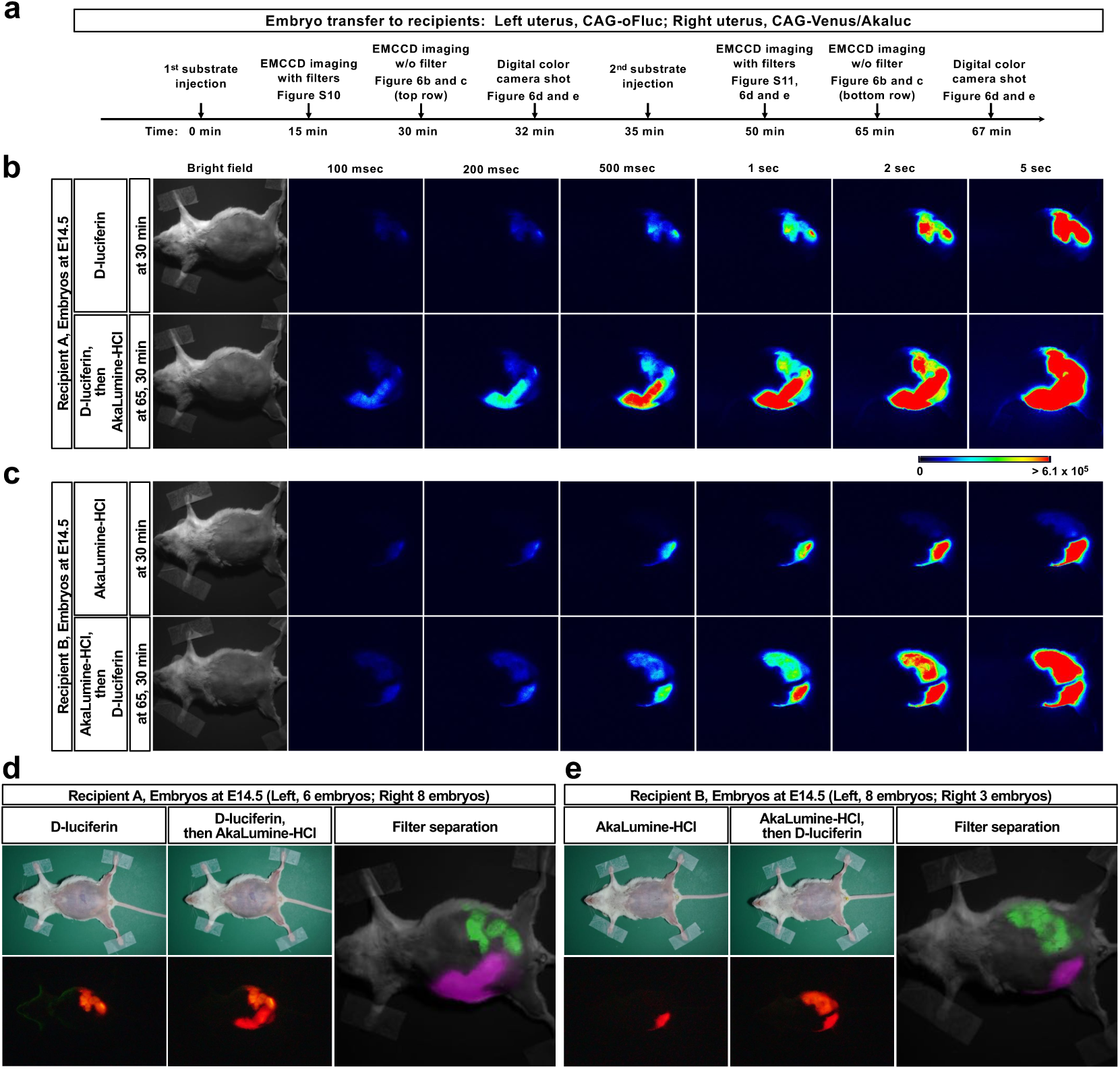
In vivo dual-color BLI of “glowing embryos” in pregnant mice Dual-color BLI of recipient mothers transplanted with CAG-oFluc and CAG-Venus/Akaluc embryos into the left and right uteri, respectively. (a) Timing of substrate injection and imaging. (b and c) In vivo BLI without optical filters. (b) Recipient A at 30 min after the injection of the first substrate (top, D-luciferin, 100 mM D-luciferin, 10 μl/gbw) and at 30 min after the injection of the second substrate (bottom, AkaLumine-HCl, 3 mM AkaLumine-HCl, 5 μl/gbw). The first and second substrates were injected sequentially at 35-min intervals; therefore, the images at the bottom were acquired at 65 min after the injection of the first substrate. Number of embryos in Recipient A: 6 CAG-oFluc and 8 CAG-Venus/Akaluc embryos in the left and right uteri, respectively. (c) Recipient B, in which substrates were injected in the opposite order to that in Recipient A. Number of embryos in Recipient B: 8 CAG-oFluc and 3 CAG-Venus/Akaluc embryos in the left and right uteri, respectively. In Recipient B, AkaLumine-HCl was used at a half dose compared with Recipient A. (d and e) Images acquired using a digital color camera after the injection of the first (left) or second (middle) substrate. Filter segregation of oFluc and Akaluc (right). After the injection of the second substrate, images were captured by the EM-CCD camera using one of two optical filters (see Supplementary Figure S11). The images were pseudo-colored and merged.

The experimental scheme is illustrated in Figure 6a. After either D-luciferin or AkaLumine-HCl was injected as the first substrate, the side with fetuses with the correct luciferase/luciferin pair began producing a substantial signal, by contrast, the other side did not (Figure 6b and c, top row), indicating that minimal substrate cross-reactivity occurred in single mice. After injection of the second substrate, we observed that all the transplanted fetuses could glow at both sides comparably in either case (Figure 6b and c, bottom row).

As the images described above were acquired without any optical filter, no information regarding light wavelengths was obtained. Nevertheless, the difference in emitting colors could be appreciated based on images acquired using the digital color camera (Figure 6d and e, left half). To separate the oFluc and Akaluc signals spectrally, we used two band-pass (BP) filters, i.e., 565 ± 40 BP and 730 ± 45 BP (Supplementary Figure S9). Although the oFluc/D-luciferin signals leaked slightly into the 730 ± 45 channel (Supplementary Figures S10 and S11, left uterus imaged with the 730 ± 45 BP filter), the signals of the respective luciferases were effectively separated and sufficiently strong to compose clear merged images (Figure 6d and e, right half).

## Discussion

In the present study, we generated and characterized reporter mouse strains for the new BLI system using the highly bright luciferases, oFluc and Akaluc. Akaluc is 100–1000-fold more sensitive for deep-tissue imaging than the conventional BLI system ^20^. oFluc is a novel luciferase that produces at least 10-fold more intense light than the commonly used luciferase (Luc2) in vitro ^23^. In the reporter mouse strains, Cre-dependent reporter constructs were knocked into the *ROSA26* locus and driven by the strong *CAG* promoter.

After crossing these mice with a Cre-deleter strain, we generated “glowing mice” that emitted high-intensity light from their entire bodies and tissues. The behavior of these freely moving mice in the dark could be recorded at a video rate of 30 fps using a consumer-grade digital color camera (Supplementary Movie 1). We also successfully imaged E6.5 embryos in utero using an EM-CCD camera. However, this is probably not the limit of detection; in some cases, E5.5 mouse embryos carrying oFluc could be imaged (data not shown).

The reporter constructs used in this study were designed to be normally silent because of the transcriptional stop signal flanked by loxP sites. The expression of oFluc or Venus/Akaluc is supposed to be induced by the expression of Cre. For example, we achieved deep-tissue imaging of the pancreas, lymphoid organs, or specific brain regions by crossing with the appropriate Cre-driver strains (Figures 3 and 4). Although extremely weak BLI signals were noticed before the Cre crossing, which may be the result of transcriptional leakage, signals two orders of magnitude greater were obtained as Cre-dependent signals. Therefore, with an appropriate negative control, true BLI signals can be identified without any difficulty. Interestingly, the leaky expression of Venus/Akaluc was not detected by Venus fluorescence, suggesting low sensitivity of FP imaging in vivo. The results of an enzymatic activity assay in the tissue extracts (Figure 2) demonstrated that oFluc and Akaluc had wider dynamic ranges (on average, 1.2 × 10^5^-fold and 1.8 × 10^3^-fold, respectively) than Venus (8.5 × 10-fold).

Two factors are considered for the performance of BLI systems using intact animals. First, the biodistributions of AkaLumine-HCl and D-luciferin differ. Delivery across the BBB is well achieved with AkaLumine-HCl but not D-luciferin. Thus, the oFluc/D-luciferin system performed poorly in the brain (Figure 4), although outside the brain, its signals were as bright as or brighter than those of the Akaluc/AkaLumine-HCl system in some cases (Figure 3). Second, the longer the emission wavelength, the greater the tissue penetration of the light. Although the synthetic luciferin CycLuc1 has improved biodistribution across the BBB ^15^ and the oFluc/CycLuc1 system produced substantial signals in the brain with a 603-nm emission peak (Figure 4b), the signal strength could not match that of the Akaluc/AkaLumine-HCl system with a 650-nm emission peak. Taken together, we concluded that the AkaBLI system, composed of Akaluc and AkaLumine-HCl, is the most effective BLI system for detecting luciferase expression in the brain.

The emission peaks of oFluc and Venus/Akaluc are separated by 90 nm (Figure 1d). The yellow light produced from oFluc is not suitable for in vivo imaging because of its high absorption in the body ^4^. However, because of its very high light intensity, the oFluc signal is actually comparable with that of Akaluc in the body, with the exception of the brain (Figures 3 and 4). These two luciferases have the advantages of high luminosity, sufficiently separated emission peaks, and low cross-reactivity between D-luciferin and AkaLumine-HCl (Figure 3 and Supplementary Figures S4–S6), rendering them excellent partners for dual-color BLI. For dual-color BLI, an optical filter for the yellow light was selected to detect the oFluc signal. Conversely, because the oFluc signal overlapped partially with the emission spectrum of Venus/Akaluc, which has a peak at 650 nm, we selected an optical filter for the longer side of the shoulder in the emission spectrum of Venus/Akaluc (Supplementary Figure S9).

Using this setup, it was possible to detect the distinct signals from the two luciferases simultaneously by injecting a single substrate followed by injecting the second substrate (Figure 6). This administration regimen allowed us to assess substrate cross-reactivity in a single mouse when injecting the first substrate, followed by dual-color imaging after injecting the second substrate. The filter setup used in this experiment effectively separated the signals of the two luciferases; nevertheless, two concerns should be noted. oFluc/D-luciferin produced marginal signals using the longer-wavelength filter (Supplementary Figures S10 and 11). However, this could be alleviated by the much stronger signal intensity of Akaluc/AkaLumine-HCl (Supplementary Figures S10 and 11). Further characterization of the spectral shifting of signals using a series of filters and the application of spectral unmixing algorithms ^38^ would improve the separation of signals from the two luciferases. Moreover, because the biodistribution of each substrate may vary within the target tissue in the body, the substrate concentrations to be administered should be carefully determined (empirically).

The availability of two luciferase reporter strains with a comparable signal brightness and distinct emission peaks should facilitate a wide variety of studies in life science fields. For example, behavior of the grafted cell can be traced by in vivo BLI for studies of regenerative medicine ^22^. The “glowing mouse” can be used as an unlimited, reliable and convenient source of luminescent tissues and cells for such transplantation studies. We learned that characteristics of luciferase substrates greatly influence the performance of BLI. Therefore, a variety of mouse strains expressing different luciferase should accelerate studies searching for better luciferin derivatives and evaluating their biodistribution in mice ^39–42^. For tissue- and cell-specific labeling using the Cre/loxP system, hundreds of Cre-driver strains have been developed to date, and their specificity has mainly been examined using fluorescent reporter mice ^24, 43^. Although fluorescent reporters are excellent for the cellular imaging of dissected tissues and sections, screening for specificity at the organismal level is a time-consuming, laborious, and often impractical process. However, the BLI systems presented here allow the rapid, unambiguous, and noninvasive evaluation of Cre specificity in the whole body. We in fact found that Emx1-Cre x Akaluc mice revealed hitherto unknown domains of the Cre recombination in several body sites other than the brain (Figure 4a). Once the target sites are narrowed down by in vivo BLI, characterization at cellular resolution can be performed using Venus fluorescence and immunohistochemical detection of luciferase. After confirming the specificity of Cre-driver strains, these strains should offer numerous research tools/materials for noninvasive in vivo BLI, to study various complex biological phenomena in live mice in healthy and diseased conditions.

## Materials and methods

### Ethical statements

All experimental protocols and husbandry for mice were approved by the Institutional Animal Care and Use Committee of RIKEN Tsukuba Branch and University of Tsukuba, and all mice were cared for and treated humanely in accordance with the Committee’s guiding principles.

### Construction of targeting vectors

Targeting vectors were constructed based on a plasmid described in a previous study ^44^. They comprised the homology arms—1.1 kb at the 5′ portion and 2.8 kb at the 3′ portion—of the *ROSA26* locus, and an insert DNA fragment including the *CAG* promoter followed by a pair of loxP sites flanking the neomycin resistance gene (*Neo*) and SV40 poly A (which were derived from pCALNL5 (RDB01862)), followed by either the luciferase derived from *Pyrocoeli matsumurai* (oFluc, RDB14359) or Venus/Akaluc (RDB15781), with the addition of a woodchuck hepatitis virus posttranscriptional regulatory element (WPRE) and bovine growth hormone ploy A at the 3′ end. The resulting insert DNA fragment of oFluc was ligated to the XbaI site of the *ROSA26* homology arm, whereas that of Venus/Akaluc was ligated to the *ROSA26* homology arm, with parts of the arms being deleted—94 bp upstream and 41 bp downstream of the XbaI site—to increase gRNA selection for CRISPR‒Cas9 genome editing.

### Generation of mouse strains

Mice were generated using the CRISPR‒Cas9 technique with zygotes derived from C57BL/6JCrl mice, as described in a previous study ^45^. The target sequences of gRNAs in the *ROSA26* locus were as follows: 5′–CGCCCATCTTCTAGAAAGAC–3′ for oFluc and 5′–TGGCTTCTGAGGACCGCCCT–3′ for Venus/Akaluc. Cas9, gRNA, and the targeting vector were microinjected in the form of nonlinearized plasmid DNAs, and then pups born after zygote implantation into the oviducts of foster mothers were screened by PCR. An initial screening was carried out to confirm homologous recombination targeting the *ROSA26* locus using PCR primers that annealed to the genomic region external to the homology arms and the insert DNA fragment corresponding to *CAG* or WPRE for the 5′ and 3′ regions, respectively. The integration of the plasmid backbone of the targeting vector was also tested using PCR primers that detected the ampicillin-resistance gene. These primer sets are listed in Table 1. Founder mice were selected based on 5′ and 3′ homologous recombination at the *ROSA26* locus, regardless of the presence of the plasmid backbone. The founder mice were further bred with C57BL/6JCrl mice.

### Evaluation of targeted knock-in in germ line-transmitted mouse lines

The genotypes of N1 pups from founder mice were tested using the PCR primers described in the previous section. To assess the genomic structure, further PCR was conducted using additional primers that could amplify the plasmid backbone used in the targeting vectors. In addition, the copy number of the insert DNA fragment integrated into the mouse genome was determined by quantitative PCR (qPCR) with separate amplification of two components—*Neo* and WPRE; their copy number was compared with a standard genomic DNA (RBRC04874) that was previously confirmed to represent the integration of 1 copy into the *ROSA26* locus ^44^. Targeted knock-in was evaluated in mice after the CAG-Cre cross, as described above. Furthermore, the absence of floxed *Neo* was tested using two primer sets, one for *Neo* and the other for the regions upstream and downstream of *Neo*. The primers used in this section are listed in Table 1.

### Animal breeding

All mice were provided with commercial laboratory mouse diet and water ad libitum, and were housed under lighting conditions (light on from 8:00 to 20:00) and specific pathogen-free conditions. The Cre-driver mice used in this study were as follows: C57BL/6-Tg(CAG-cre)13Miya (RBRC09807, CAG-Cre), B6.129P2-Emx1<tm1.1(cre)Ito>/ItoRbrc (RBRC01345, Emx1-Cre), C57BL/6J-Tg(Slc32a1-cre)65Kzy (RBRC10606, Vgat-Cre), B6;Cg-Pdx1<tm1(cre)Yasu> (RBRC10170, Pdx1-Cre), and B6.Cg-Tg(Lck-cre)1Jtak (RBRC04738, Lck-Cre). All of these Cre-driver mice were obtained from RIKEN BRC. In general, male Cre-driver mice were bred with female reporter mice to generate compound Cre- and reporter-positive mice. B6.Cg-c/c Hr<HR> (RBRC05798) mice were used to introduce the albino and/or hairless phenotype only when bioluminescent imaging was conducted in freely behaving mice. All mouse strains and their genotyping protocols are available from RIKEN BRC at https://mus.brc.riken.jp/en/. For in vivo imaging of pregnant females carrying E6.5 embryos, BALB/c mice were purchased from CLEA Japan and crossed with CAG-oFluc or CAG-Venus/Akaluc mice. Some of their pups were imaged at postnatal day 4.

### Evaluation of luciferase enzymatic activity and of the fluorescence intensity of crude tissue extracts

Mice were sacrificed via cervical dislocation and their tissues were immediately dissected, snap-frozen in liquid nitrogen, and stored at –80 °C until use. The frozen tissues (50–250 mg) were lysed with a 5× volume of lysis buffer containing 25 mM Tris/phosphate, 4 mM EGTA, 1% Triton X-100, 10% glycerol, 2 mM dithiothreitol, and an EDTA-free protease inhibitor cocktail (Roche, Tokyo, Japan), followed by homogenization with zirconia beads in a refrigerated Micro Smash MS-100R instrument (TOMY, Tokyo, Japan) for 30 s twice. After brief centrifugation, the supernatants were transferred to clean tubes and their protein concentrations were measured using a BCA Protein Assay Kit (Pierce Thermo Scientific, MA, USA), according to the manufacturer’s instructions. Subsequently, 20 μl of the supernatant was mixed with 180 μl of the assay buffer containing 25 mM Tris/phosphate, 20 mM MgSO_4_, 4 mM EGTA, 2 mM ATP, 1 mM dithiothreitol, and 1 mM substrate in a black-walled 96-well plate (Thermo Fisher Scientific, MA, USA), and then immediately measured on a luminometer (BioTek Synergy HTX, Agilent Technologies, CA, USA). The values of bioluminescence recorded at 5 min after mixing were compared among genotypes. Stock solutions of the three substrates, i.e., D-luciferin (Cayman Chemical Company, MI, USA), AkaLumine-HCl (Wako, Osaka, Japan), and CycLuc1 (MedChemExpress, NJ, USA), were dissolved in PBS, distilled water, and PBS at final concentrations of 100, 100, and 5 mM, respectively. In addition, the fluorescence intensity of Venus/Akaluc was measured using a BioTek Synergy HTX reader.

### Measurement of the emission spectra of recombinant oFluc and Venus/Akaluc

The cDNA of either oFluc (RDB14359 from RIKEN BRC) or Venus/Akaluc ^20^ was inserted in the multiple cloning site of pRSET-B using BamHI and EcoRI (Thermo Fisher Scientific, MA, USA). The luciferase protein expressed in NiCo21(DE3) competent *Escherichia coli* (New England Biolabs, MA, USA) was purified using a Ni-NTA agarose resin column (Qiagen, Venlo, Limburg, Netherlands) and a chitin resin column (New England Biolabs, MA, USA). Protein concentrations were measured using a Bradford Protein Assay (BIO-RAD, CA, USA). Emission spectra were determined using a LumiFl-Spectrocapture AB-1850 instrument (Atto, Tokyo, Japan) at 37 °C in a solution of 0.1 M citrate phosphate buffer (pH 7.0) containing the substrate (1 mM D-luciferin or 10 μM AkaLumine), 1 mM ATP, 2 mM MgSO_4_, and each of the purified luciferases (10 ng/μl in 100 μl reaction volume).

### Measurement of the Michaelis–Menten constant (Km) values

Km for oFluc or Venus/Akaluc for D-luciferin or AkaLumine were determined with a luminometer, Nivo S (PerkinElmer, MA, USA). Luminescence intensity was measured in 0.1 M citric acid / 0.1 M Na_2_HPO_4_ buffer (pH 7.0) containing 2 μg/μl of partially purified luciferase, 1 mM ATP, 2 mM MgSO_4_, and luciferin (D-luciferin 0-990 μM, AkaLumine 0-208 μM). The time course of light emission was measured for 10 s. The Km was estimated by curve fitting against the Michaelis–Menten equation.

### Ex vivo BLI

Tissues were isolated immediately after the cervical dislocation of mice and the dissected tissues were incubated in PBS containing 1 mM substrate for 5 min. Subsequently, BLI signals were captured with VISQUE InVivo Smart-LF (Vieworks, Gyeonggi-do, Korea) and analyzed using the accompanying software, CleVue. The fluorescence signals were also imaged with InVivo Smart-LF. Dissected brains and testes (Figure 5 and Supplementary Figure S3) were imaged with an ImagEM 9100-13 (Hamamatsu Photonics, Shizuoka, Japan) and a lens (#903018, AstroScope, NY, USA) set up in a dark chamber, and the acquired images were analyzed using CellSense software (Olympus,Tokyo, Japan). The signal intensities captured by ImagEM 9100-13 were calibrated using KoshiUni (Atto, Tokyo, Japan), to determine its sensitivity, and are presented as the absolute photon number per second and cm^2^.

### In vivo BLI

Before the imaging session, any hair covering the region of interest was removed using a shaver followed by depilatory cream. Mice were anesthetized with a combination of anesthetics, as follows: 0.3 mg/kg medetomidine, 4.0 mg/kg midazolam, and 5.0 mg/kg butorphanol. Bioluminescence images were acquired with an ImagEM 9100-13 (Hamamatsu Photonics, Shizuoka, Japan) and a lens (#903018, AstroScope, NY, USA) set up in a dark chamber. In each imaging session, one of three substrates was injected intraperitoneally at the dose indicated in the Results section in proportion to the grams of body weight (gbw) of mice, and image acquisition generally started 10 min after substrate administration at multiple exposure times. The images were analyzed using CellSense software. Movies of freely behaving mice were recorded using a digital color camera, α7SII, with the following settings: ISO, 102400; f2.9 lens (Sony, Tokyo, Japan) in a dark room. For successive imaging sessions using multiple substrates, each imaging session was separated by at least 24 h and the absence of the luminescence signal emitted from the previous imaging session was confirmed at the beginning of each new imaging session.

Data from images acquired by an ImagEM 9100-13 were quantified using ImageJ software (NIH) based on the calibration data provided by KoshiUni (Atto, Tokyo, Japan). Fluorescence images were also acquired using a VISQUE InVivo Smart-LF or Keyence GFP-lighting system (VB-L12, Keyence, Osaka, Japan) equipped with a CCD camera (DFC450C, Leica Microsystems, Wetzlar, Germany).

### Histology

The dissected tissues were fixed with 4% paraformaldehyde in 0.1 M phosphate buffer overnight, and then embedded in OCT compound (Sakura, Tokyo, Japan). The tissues were further treated with 30% sucrose overnight. Cryostat sections were prepared for native fluorescence imaging and anti-Luc2 (MBL, Tokyo, Japan) immunohistochemistry (IHC). For anti-Luc2 IHC, the tissue sections were first blocked with PBS containing 3% normal donkey serum and 0.3% Triton-X100, and then incubated with a rabbit anti-Luc2 antibody (1000× dilution) overnight at 18 °C. After washing three times with PBS, the sections were further incubated with ImmPRESS (Vectorlabs, CA, USA) for 2 h at 24 °C. Signals were visualized with TSA-Cy3 (PerkinElmer, MA, USA). All sections were counterstained with DAPI (Thermo Fisher Scientific, MA, USA).

### In vivo dual-color BLI

Heterozygous fertilized eggs were prepared via in vitro fertilization using sperm from homozygous males of either the CAG-oFluc or CAG-Venus/Akaluc strains and eggs of C57BL/6. In vivo dual-color BLI was conducted with two ICR recipients (CLEA Japan) in which heterozygous CAG-oFluc embryos had been transferred into the left uteri and heterozygous CAG-Vensu/Akaluc embryos into the right uteri. Bioluminescence images were acquired with an ImagEM 9100-13 (Hamamatsu Photonics, Shizuoka, Japan) and a lens (HF25HA-1B, Fujinon, Saitama, Japan), without any filter, which were set up in a dark chamber. Images were further acquired using two band-pass filters (565 ± 40 BP and 730 ± 45 BP from Omega Optical, VT, USA) attached to the lens sequentially to separate the signals from oFluc and Venus/Akaluc. Substrate dosage and the timing of their injection and imaging are described in the Results section and Figure 6a.

### Statistical analysis

All data are presented as the mean ± standard error of the mean (SEM). Data between groups were compared using a Student’s *t* test. The null hypothesis was rejected at the *P* < 0.05 level.

## Acknowledgements

We thank Drs. J. Miyazaki, S. Itohara, Y. Yamada, J. Takeda, and K. Ochiai for their generous deposition of mouse strains in the RIKEN BRC and for allowing their use in this study. We also thank the members of the Experimental Animal Division of RIKEN BRC, especially M. Kadota, Y. Yamashita, and K. Otaka, for their excellent technical assistance, and Dr. M. Sugimoto for his help with embryo imaging. This work was supported by JSPS KAKENHI Grant No. JP26890026 to T.N., and in part by JST Grant No. JPMJPF1234 to A.Y. The project was also supported in part by the RIKEN BRC director’s discretionary fund.

## Competing interests

The authors have declared that no competing interests exist.

## Author contributions

T.N. conceived, designed, and performed all the experiments (except for the emission spectra of recombinant proteins), analyzed all data, and drafted the manuscript.

K.O. conceived, designed, and performed the experiment to measure the emission spectra.

S.I. conceived the project and provided critical input on this study.

T.S. conceived and supported the entire project, especially regarding the imaging setup.

S.M-I. conducted the initial screening of founder mice.

K.N. conducted the initial screening of founder mice.

S.M. generated founder mice.

F.S. generated founder mice.

A.Y. managed the animal facility and secured funding.

A.M. conceived the project, provided critical input on this study and edited the manuscript.

K.A. conceived and supervised the project, and drafted the manuscript.

## Notes

### Competing Interest Statement

The authors have declared no competing interest.

