## Supplementary for "Development of two mouse strains conditionally expressing bright luciferases with distinct emission spectra as new tools for in vivo imaging"

### Supplementary figures and table

#### Figure S1

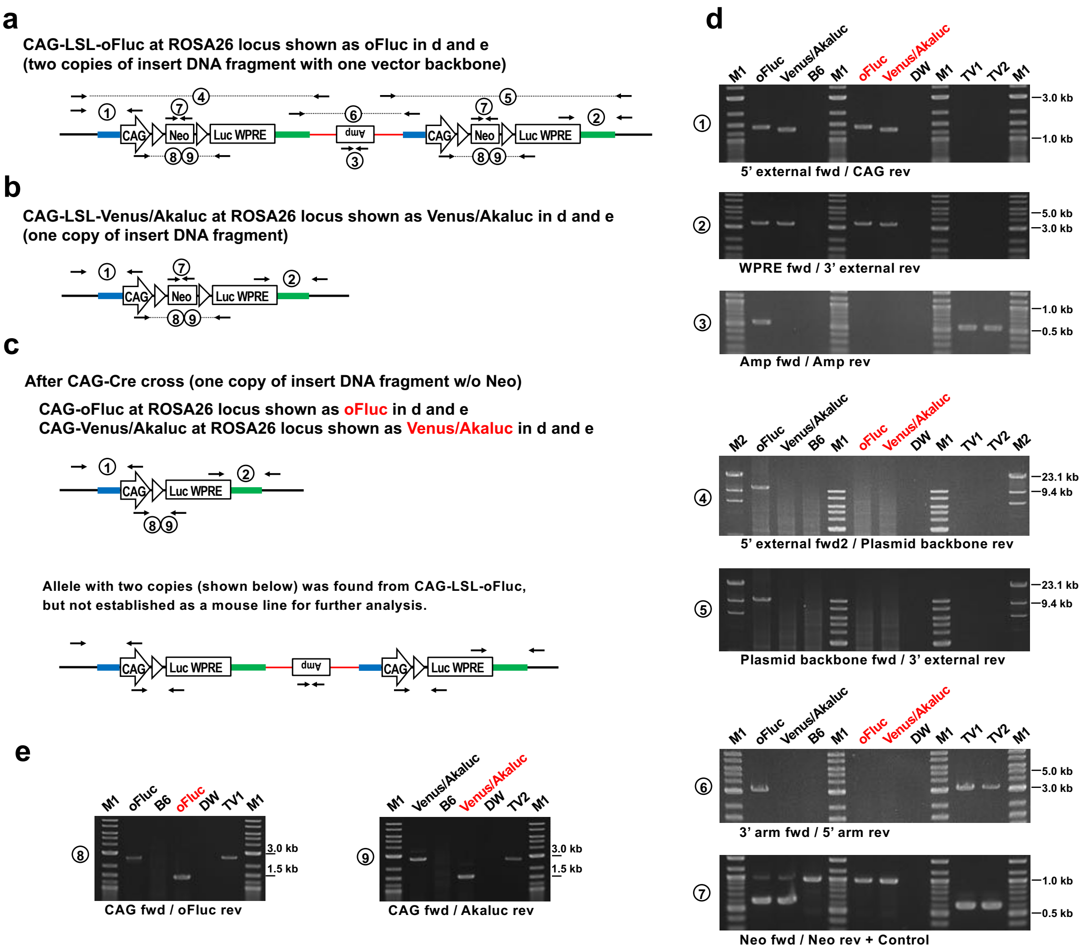


**Figure S1.** PCR confirmation of targeted knock-in at the *ROSA26* locus in luciferase reporter mice

(a) CAG-LSL-oFluc mice carried two copies of the insert DNA fragment (Figure 1a) with a vector backbone at the *ROSA26* locus. (b) CAG-Venus/Akaluc mice carried one copy of the insert DNA fragment at the *ROSA26* locus. (c) After the CAG-Cre cross, both CAG-oFluc and CAG-Venus/Akaluc mice had one copy of the insert DNA fragment with deletion of the *Neo* cassette. The locations of the PCR primers are indicated by numbers corresponding to the gel images indicated in (d) and (e). (d) Confirmation of the targeted knock-in at the *ROSA26* locus by PCR. The PCR primer sets used in each panel are indicated by numbers in (a) to (c). oFluc in black (CAG-LSL-oFluc), Venus/Akaluc in black (CAG-LSL-Venus/Akaluc), B6 (C57BL/6), oFluc in red (CAG-oFluc), Venus/Akaluc in red (CAG-Venus/Akaluc), DW (distilled water, negative control), TV1 (targeting vector plasmid for oFluc), and TV2 (targeting vector plasmid for Venus/Akaluc). M1 and M2 are DNA size markers (i.e., Quick-Load 1 kb Plus DNA Ladder and λDNA-HindIII Digest, respectively (NEB, Japan)). (e) Confirmation of Cre-mediated recombination by PCR. The samples and PCR primers are the same as those described in (d).

#### Figure S2

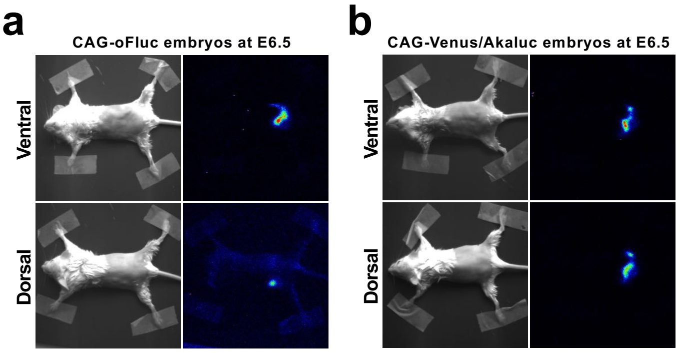


**Figure S2.** In vivo BLI of pregnant wild-type mothers carrying CAG-oFluc (a) and CAG-Venus/Akaluc (b) embryos at 6.5 days after conception. The hair on the lower abdomen and back of the mice was shaved.

### Figure S3

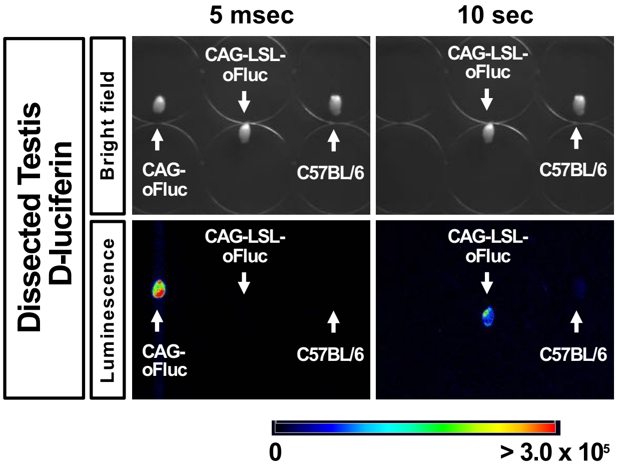


**Figure S3.** Ex vivo imaging of dissected testes incubated with D-luciferin (1 mM). For longer exposure, the testis dissected from a CAG-oFluc mouse was removed during BLI because its bright luminescence lit up other testes, thus leading to false-positive signals.

#### Figure S4

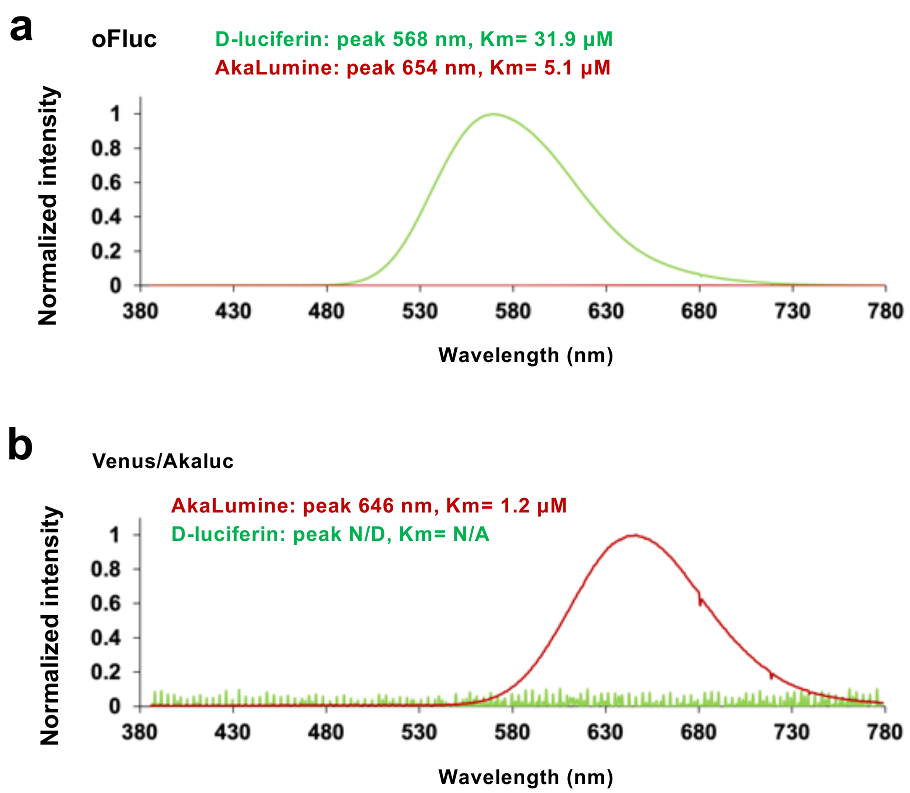


**Figure S4.** Emission spectra of the recombinant proteins of oFluc (a) and Venus/Akaluc (b), where D-luciferin or AkaLumine was used as their substrate. The intensity was normalized by dividing the measured intensity along each wavelength by the peak intensity of the matched luciferase/luciferin pairs, i.e., the peak intensity of D-luciferin at 568 nm for oFluc (a) and the peak intensity of AkaLumine at 646 nm for Venus/Akaluc (b). Emission spectra of the matched luciferase/luciferin pairs in this figure were reused from Figure 1d.

**Figure S5**

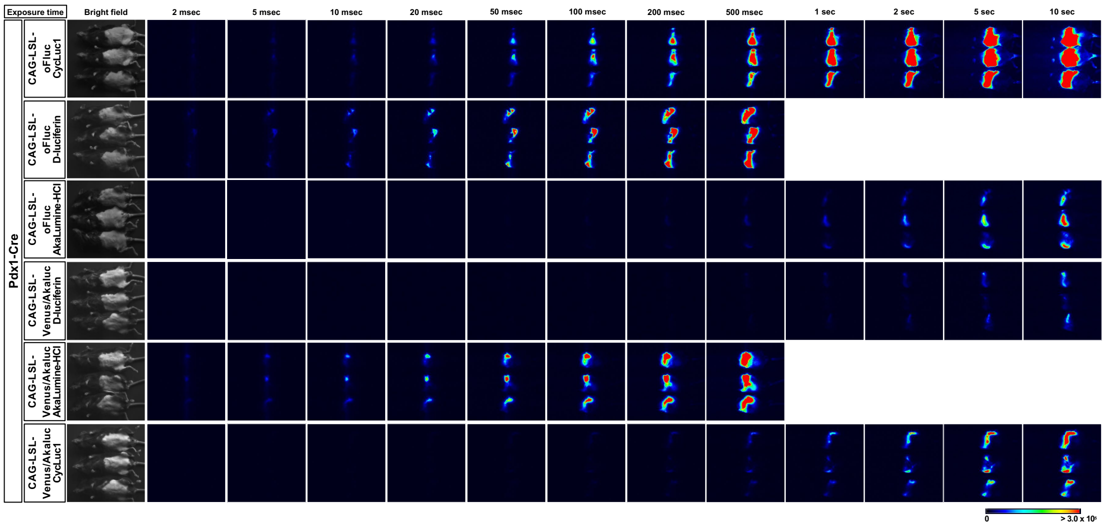


**Figure S5.** In vivo BLI of oFluc and Akaluc in the pancreas by crossing Pdx1-Cre mice with CAG-LSL-oFluc or CAG-LSL-Venus/Akaluc mice. Images were acquired using multiple exposure times after the administration of one of the three substrates. Data from these images were quantified as reported in Figure 3a. Scale, photon/s/cm$^2$.

**Figure S6**

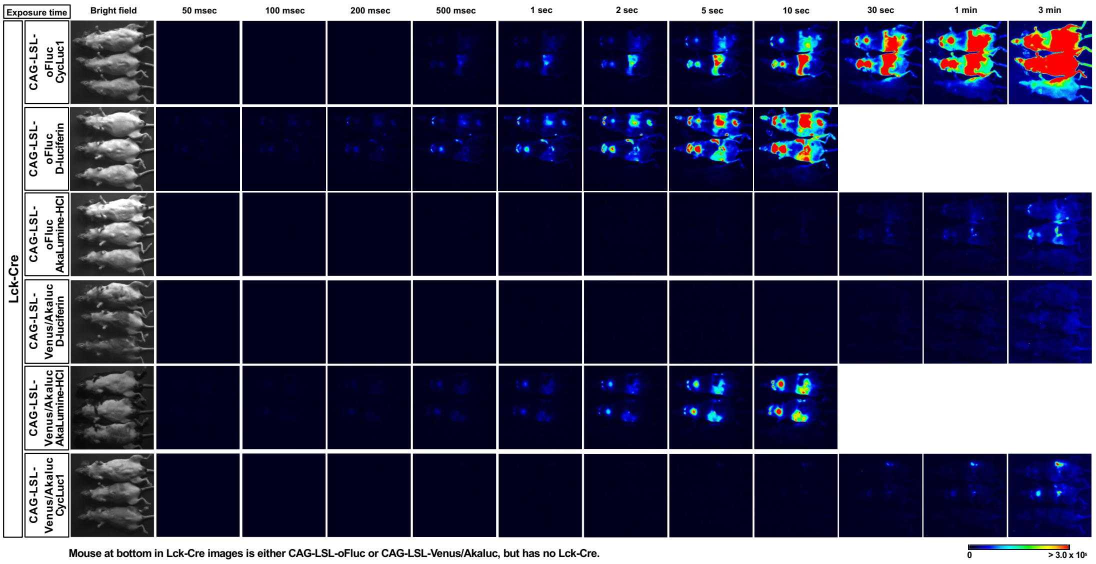


**Figure S6.** In vivo BLI of oFluc and Akaluc in T cells by crossing Lck-Cre mice with CAG-LSL-oFluc or CAG-LSL-Venus/Akaluc mice. Images were acquired using multiple exposure times after the administration of one of the three substrates. Data from these images were quantified as depicted in Figure 3b. Scale, photon/s/cm$^2$.

**Figure S7**

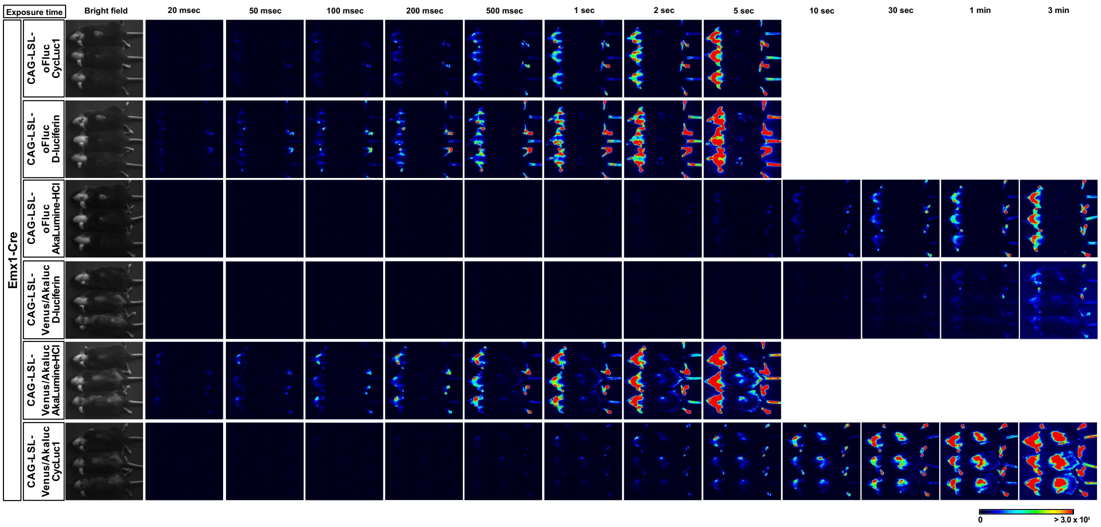


**Figure S7.** In vivo BLI for oFluc and Akaluc in mice that were double positive for Emx1-Cre and the reporters. Images were acquired using multiple exposure times after the administration of one of the three substrates. Data from these images were quantified as depicted in Figure 4a. Scale, photon/s/cm$^2$.

**Figure S8**

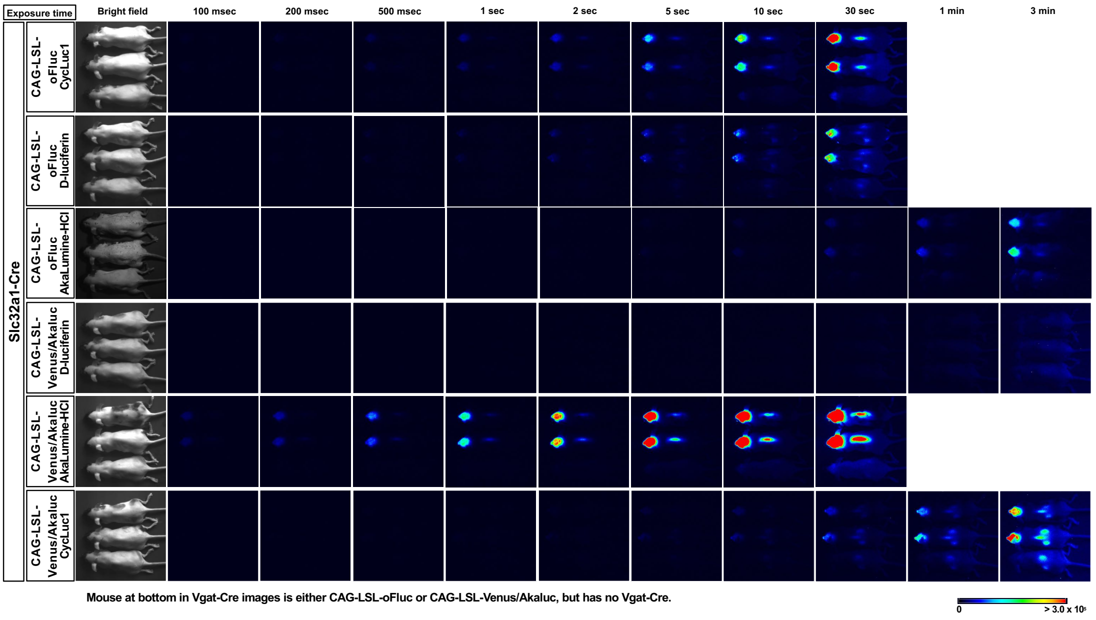


**Figure S8.** In vivo BLI for oFluc and Akaluc in mice that were double positive for Vgat-Cre and the reporters. Images were acquired using multiple exposure times after the administration of one of the three substrates. Data from these images were quantified as depicted in Figure 4b. Scale, photon/s/cm$^2$.

#### Figure S9

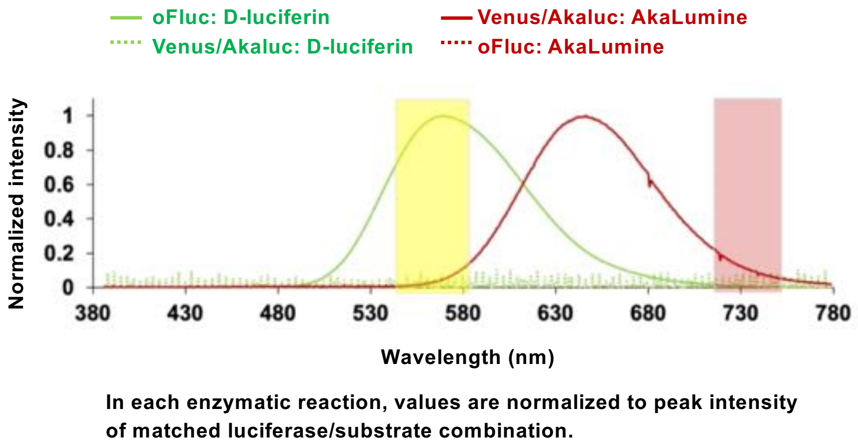


**Figure S9.** Emission spectra of the recombinant proteins of oFluc and Venus/Akaluc. Solid lines, matched luciferase/luciferin pairs; dashed lines, mismatched luciferase/luciferin pairs. The yellow and red boxes indicate the two wavelength ranges that can pass through the filters used in the in vivo dual-color BLI, as depicted in Figure 6 and Supplementary Figures S10 and S11. Yellow box, 565 ± 40 BP filter; red box, 730 ± 45 BP filter. This figure was generated by combining Figure S4a and S4b.

**Figure S10**

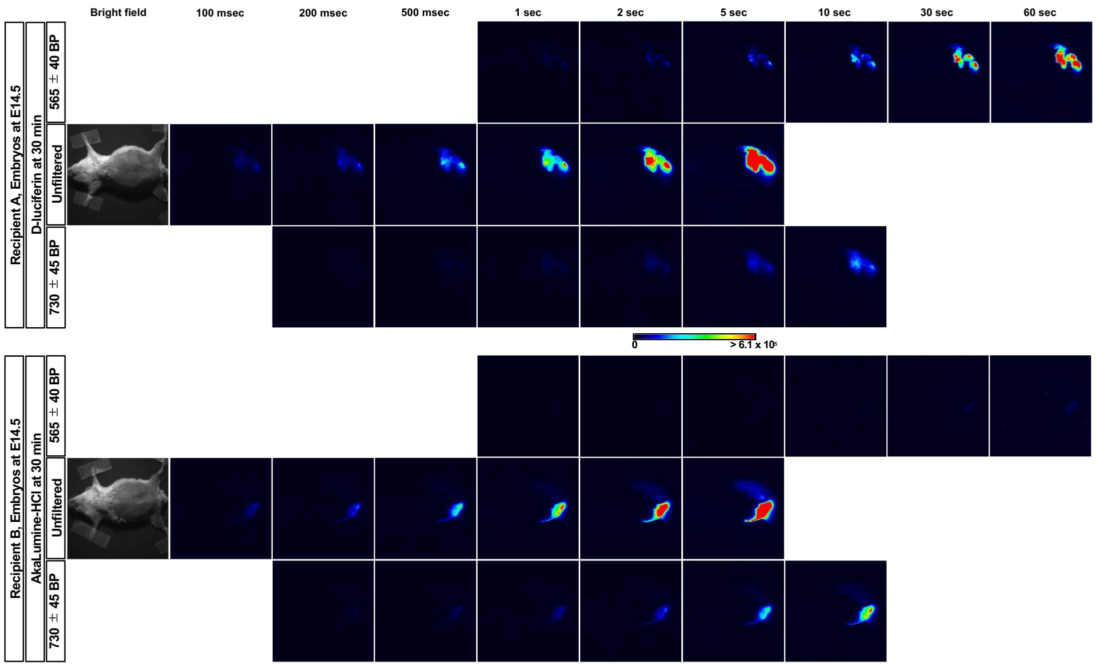


**Figure S10.** Filter segregation of oFluc and Akaluc after injecting the first substrate, as depicted in Figure 6.

**Figure S11**

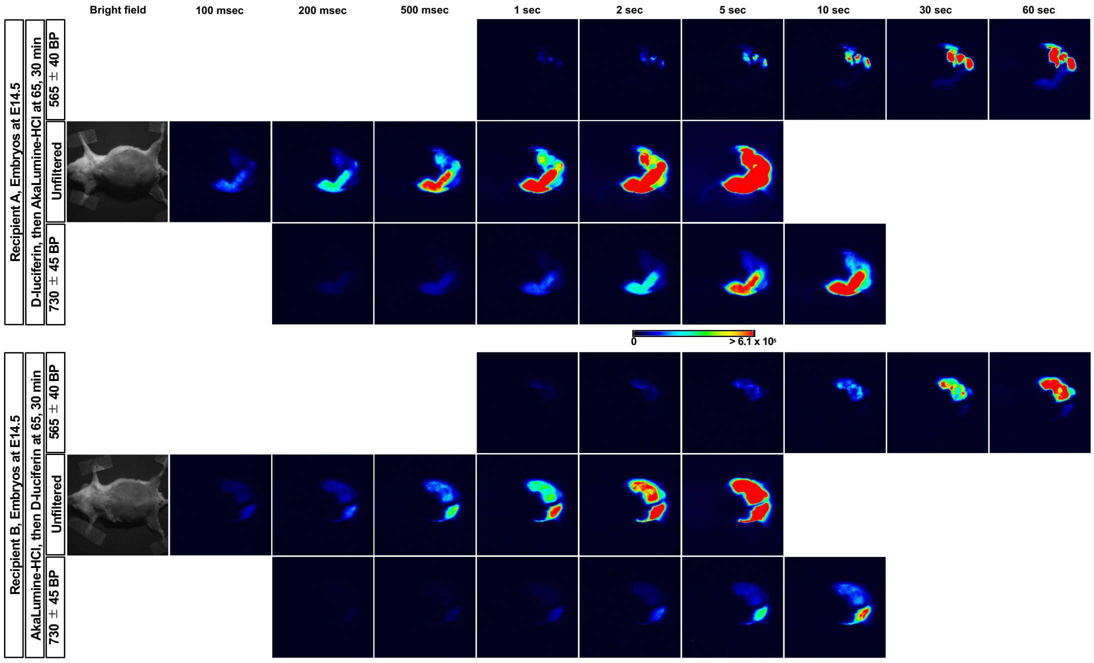


**Figure S11.** Filter segregation of oFluc and Akaluc after injecting the second substrate, as depicted in Figure 6.

**Table S1**

| PCR test | Primer name | Primer sequence | CAG-LSL-oFluc | CAG-LSL-Venus/Akaluc | B6 (C57BL/6) | CAG-oFluc | CAG-Venus/Akaluc | DW (Distilled water) | TV1 | TV2 |
|---|---|---|---|---|---|---|---|---|---|---|
| 1 | 5’ external fwd | GCCCACCGCCCCACACTTAT | 1452 bp | 1350 bp | N/A | 1452 bp | 1350 bp | N/A | N/A | N/A |
| | CAG rev | ACGTCAATGGAAAGTCCCTATTGGCGTTAC | | | | | | | | |
| 2 | WPRE fwd | GTCCCTTCGGCCCTCAATCCAGCG | 3541 bp | 3472 bp | N/A | 3541 bp | 3472 bp | N/A | N/A | N/A |
| | 3’ external rev | GCCTATTCACCGGAGAAATCCATGTCCCA | | | | | | | | |
| 3 | Amp fwd | TTGCCGGGAAGCTAGAGTAA | 560 bp | N/A | N/A | N/A | N/A | N/A | 560 bp | 560 bp |
| | Amp rev | TTTGCCTTCCTGTTTTTGCT | | | | | | | | |
| 4 | 5’ external fwd2 | GCCCACCGCCCCACACTTATTGG | 9916 bp | N/A | N/A | N/A | N/A | N/A | N/A | N/A |
| | Plasmid backbone rev | TTGTGAGCGGATAACAATTTCACACAGG | | | | | | | | |
| 5 | Plasmid backbone fwd | CGCTTACAATTTCCATTCGCCATTCAGG | 10060 bp | N/A | N/A | N/A | N/A | N/A | N/A | N/A |
| | 3’ external rev | GCCTATTCACCGGAGAAATCCATGTCCCA | | | | | | | | |
| 6 | 3’ arm fwd | CTGCAACAAGATATGTAGACTAAAGTTCTGCC | 3348 bp | N/A | N/A | N/A | N/A | N/A | 3348 bp | 3351 bp |
| | 5’ arm rev | GTCCGGTGGAGACTTTTCCGCTCC | | | | | | | | |
| 7 | Neo fwd | TGGATTGCACGCAGGTTCTC | 618 bp | 618 bp | N/A | N/A | N/A | N/A | 618 bp | 618 bp |
| | Neo rev | CGGCCACAGTCGATGAATCC | | | | | | | | |
| | Control fwd | TACCCAGTAGAGTCTATAGG | 1007 bp | 1007 bp | 1007 bp | 1007 bp | 1007 bp | N/A | N/A | N/A |
| | Control rev | AAATCCAAGTGAGGGTACGC | | | | | | | | |
| 8 | CAG fwd | GCCTTCTTCTTTTTCCTACAGCTCC | 2629 bp | N/T | N/A | 1391 bp | N/T | N/A | 2629 bp | N/T |
| | oFluc rev | TGTCGATCAGGGCATTGGTGGCC | | | | | | | | |
| 9 | CAG fwd | GCCTTCTTCTTTTTCCTACAGCTCC | N/T | 2755 bp | N/A | N/T | 1550 bp | N/A | N/T | 2755 bp |
| | Akaluc rev | GTGCGGTGCGGTAGGGCTACGCC | | | | | | | | |
